# Myosin V regulates spatial localization of different forms of neurotransmitter release in central synapses

**DOI:** 10.1101/2020.12.17.423194

**Authors:** Dario Maschi, Michael W. Gramlich, Vitaly A. Klyachko

## Abstract

Synaptic active zone (AZ) contains multiple specialized release sites for vesicle fusion. The utilization of release sites is regulated to determine spatiotemporal organization of the two main forms of synchronous release, uni-vesicluar (UVR) and multi-vesicular (MVR). We previously found that the vesicle-associated molecular motor myosin V regulates temporal utilization of release sites by controlling vesicle anchoring at release sites (Maschi et al, 2018). Here we show that acute inhibition of myosin V shifts preferential location of vesicle docking away from AZ center towards periphery, and results in a corresponding spatial shift in utilization of release sites during UVR. Similarly, inhibition of myosin V also reduces preferential utilization of central release sites during MVR, leading to more spatially distributed and temporally uniform MVR that occurs farther away from the AZ center. Thus myosin V regulates both temporal and spatial utilization of release sites during two main forms of synchronous release.

## INTRODUCTION

Neurotransmitter release is governed by the fusion of synaptic vesicles at specialized release sites at the synaptic active zone (AZ). The number, spatial distribution and temporal utilization of release sites are thought to play important roles in regulating synaptic transmission (Neher, 2010). Nanoscale imaging techniques have recently made it possible to detect individual vesicle release events in central synapses revealing the presence of multiple discrete release sites within the individual AZ. The number of release sites vary widely across the synapse population with estimates ranging from 2-18 per AZ (Tang et al., 2016; Maschi and Klyachko, 2017; Sakamoto et al., 2018). These release sites are distributed throughout the AZ with the nearest-neighbor distances of ~80-100nm, and co-localize with clusters of pre-synaptic docking factors (Tang et al., 2016). Importantly, release site usage is not uniform across the AZ, but rather forms a gradient decreasing from the AZ center to periphery with a ~4-fold difference in basal release probability between most central and most peripheral release sites (Maschi and Klyachko, 2020). Release site usage is also dynamically regulated: vesicle release preferentially occurs at more central release sites during low activity, but shifts away from AZ center towards more peripheral release sites during high-frequency stimulation (Maschi and Klyachko, 2017).

In addition to uni-vesicle release (UVR) when a single vesicle fuses in response to an action potential, a multi-vesicular release (MVR) is also prominent in many central synapses (Auger et al., 1998; Auger and Marty, 2000; Chanaday and Kavalali, 2018; Christie and Jahr, 2006; Huang et al., 2010; Korn et al., 1994; Leitz and Kavalali, 2011, 2014; Malagon et al., 2016; Rudolph et al., 2011; Singer et al., 2004; Tong and Jahr, 1994; Wadiche and Jahr, 2001). This form of synchronous release involves fusion of two or more vesicles in response to a single action potential in the same synapse and has been suggested to serve a wide range of functions including enhancing synaptic reliability, controlling synaptic integration and induction of several forms of plasticity (Rudolph et al., 2015). We recently found that MVR events exhibit spatial and temporal patterns of organization which are determined by the gradient of release site properties across the individual AZs. MVR events are also often not perfectly synchronized and are spatially organized with the first of the two events comprising MVR located closer to the AZ center (Maschi and Klyachko, 2020).

Thus the spatiotemporal organization of the two major forms of synchronous release, UVR and MVR, are both determined by the distribution of release site properties across individual AZs. Yet the mechanisms controlling the heterogeneity and utilization of release sites at the AZ in central synapses are only beginning to emerge. Recent studies suggest that release site refilling and utilization requires actin and myosins (Miki et al., 2016; Miki et al., 2018). Among actin-dependent motors, myosin V is the principle motor known to be associated with presynaptic vesicles in central neurons (Takamori et al., 2006). We recently found that acutely inhibiting myosin V markedly reduces the probability of release site reuse, and causes a profound vesicle anchoring/docking defect (Maschi et al., 2018). This is consistent with EM observations of reduced number of docked vesicles in neuroendocrine cells upon myosin V inhibition (Desnos et al., 2007). Our single-vesicle tracking measurements revealed that vesicles undergo cycles of docking and undocking at the AZ and that myosin V controls vesicle retention at release sites in an activity-dependent manner, but not vesicle transport to the release sites (Maschi et al., 2018). This function is consistent with myosin V’s ability to interact with SNARE proteins, including syntaxin 1A and synaptobrevin, and its transition from a transporting motor to a tether in a calcium-dependent manner (Krementsov et al., 2004; Ohyama et al., 2001; Prekeris and Terrian, 1997; Watanabe et al., 2005). In addition to this role for myosin V in supporting vesicle retention at release sites, our previous results suggested that spatial distribution of release is altered by myosin V inhibition. Here we extended these studies to examine the role of myosin V in determining spatial landscape of release site usage across individual AZs and its role in regulating spatial properties of UVR and MVR.

## RESULTS

### The spatial localization of vesicle docking and release in the active zone is myosin V - dependent

Our previous studies have shown that utilization of individual release sites within an AZ forms a gradient decreasing from the AZ center to periphery (Maschi and Klyachko, 2020). In other words, more central release sites have a higher release probability (Pr) and thus are preferentially used. We also found that myosin V plays an important role in refilling of the individual release sites with vesicles (Maschi et al., 2018) and therefore it actively regulates the utilization (and thus the Pr) of release sites. To explore the role of myosin V in spatially shaping the release probability landscape across the AZs, we analyzed these datasets using three independent approaches.

First, we examined the effects of acute myosin V inhibition on the spatial distribution of individual release events in the AZ of hippocampal boutons. Briefly, our imaging approach takes advantage of a pH-sensitive indicator vGlut1-pHluorin targeted to the synaptic vesicle lumen (Balaji and Ryan, 2007; Leitz and Kavalali, 2011; Voglmaier et al., 2006) allowing detection of single vesicle release events with ~20-27 nanometer precision (Maschi and Klyachko, 2017). Single release events were evoked in individual synapses at 37°C by 1 AP stimulation at 1Hz for 120 sec (or, in some experiments, with a 10Hz train for 10 sec, repeated at 0.05 Hz with the same total recording time and number of stimuli per frequency) with a frame duration of 40ms. We previously observed that acute inhibition of myosin V with a selective agent Myovin-1 (Myo-1) or with Pentabromopseudilin (PBP) caused an increase in the average distance from release events to AZ center, particularly during high-frequency (10Hz) stimulation (Maschi et al., 2018). Indeed, such a shift in location of vesicle release upon myosin V inhibition is also evident in cumulative plots of vesicle locations, particularly during high-frequency synaptic activity (10Hz) **(Figure 1B, Table 1)**.

**Figure 1:**
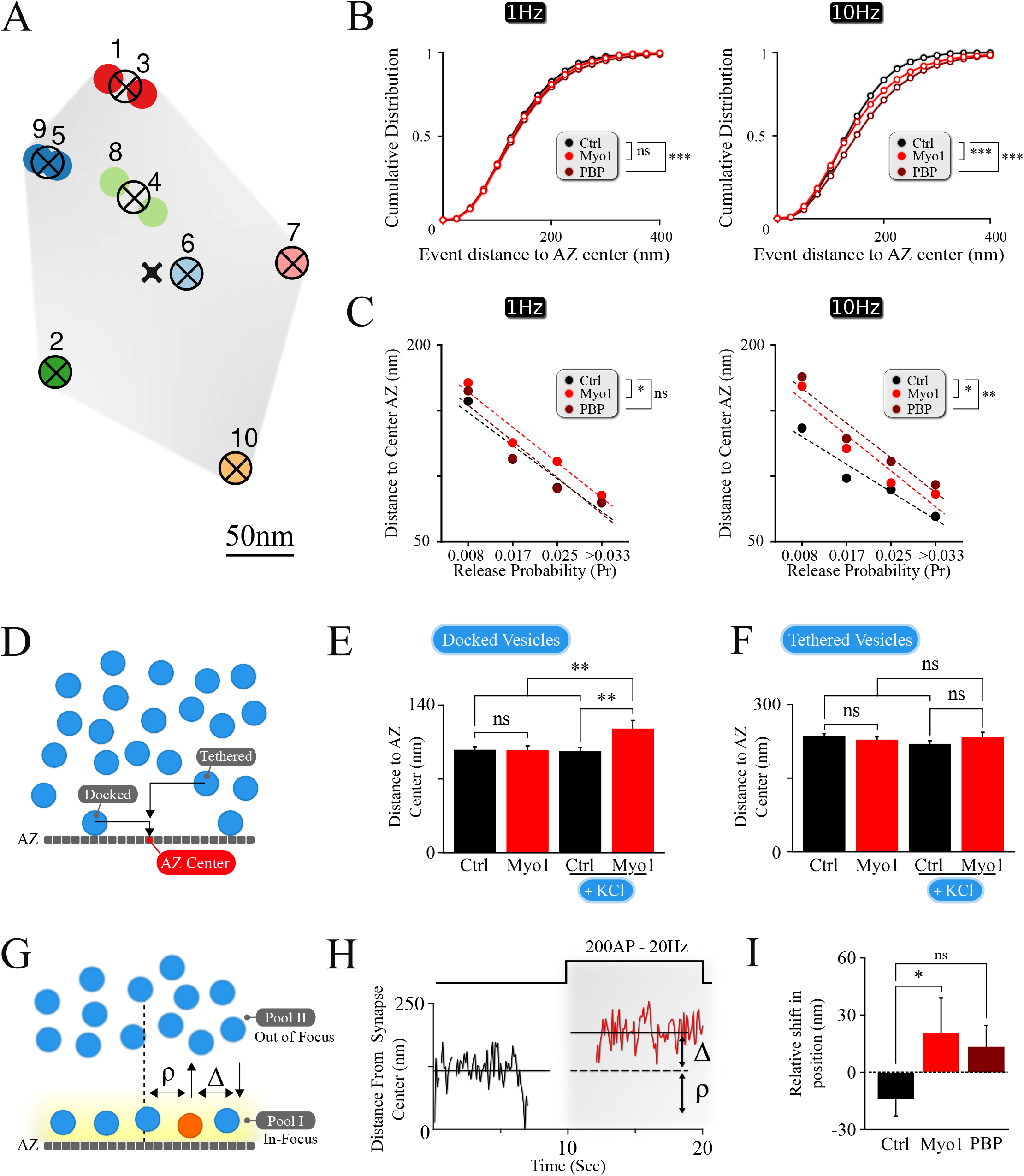
The spatial localization of vesicle docking and release is myosin V -dependent. (**A**) Sample map of release events within a single hippocampal bouton evoked by 1 Hz stimulation, with 10 fusion events and 7 release sites. Hierarchical cluster analysis was used to define release event clusters (representing individual release sites (crossed circles)) with a clustering diameter of 50 nm. Events clustered into the same release site are shown by the same color. Scale bar: 50 nm. (**B**) Effects of myosin V inhibition with Myo-1 (red) or PBP (brown) on cumulative histograms of distances from vesicle release locations to the AZ center recorded at 1Hz (left) or 10Hz (right). (**C**) Effects of myosin V inhibition with Myo-1 (red) or PBP (brown) on the average distance to the AZ center for individual release sites for measurements at 1Hz (left) or 10Hz (right), binned on the basis of their release probability. Note that errors of measurements are too small to be visible in this plot and the same data is presented as a bar-graph in **Figure1-figure supplement 1**. (**D**) Cartoon representation of the analysis of LaSEM measurements in individual hippocampal boutons in cultures depolarized (or not) by KCl application (55 mM) for 10 min in the presence or absence of Myo-1 (20 min), immediately followed by fixation. Vesicles were considered as ‘docked’ when the distance from the vesicle center to AZ was under 30 nm and ‘tethered’ when the distance was under 100 nm. (**E, F**) Effects of myosin V inhibition with Myo-1 on the localization of docked vesicles (**E**) or tethered vesicles (**F**), with or without KCl depolarization, plotted as the mean distance to AZ center (nm). (**G**) Cartoon representation of vesicle re-docking measurements using single-vesicle tracking. Vesicle disappearance/reappearance events are caused by vesicle moving out-of/back in-to the focal plane near the AZ, due to vesicle shuttling between the docking locations at the AZ and the inner vesicle pool. The relative shift in vesicle position upon re-docking was determined as a difference (Δ) of vesicle initial docking location before disappearance (ρ) and its subsequent position after re-appearance/re-docking, both measured relative to the synapse center. (**H**) Example of a single vesicle track, measured relative to the synapse center, showing a disappearances/re-appearance event. Vesicle re-appeared (red) during a 200AP, 20 Hz stimulus train farther (by Δ nm) from the initial disappearance location (ρ). (**I**) Quantification of the shift in vesicle re-appearance/re-docking location. The shift in vesicle location was determined as a difference in the exponential fits to the aggregate distributions of vesicle locations (**Figure 1-figure supplement 1C**) separated as toward synapse center versus toward periphery relative to the vesicle initial location (defined as a point of 0 shift). Errors are residual sum of squares from the exponential fits. Two-sample t-test (C, E, F) or two-sample KS-test of cumulative distributions (B, I). *p<0.05, **p<0.01, ns - not significant.

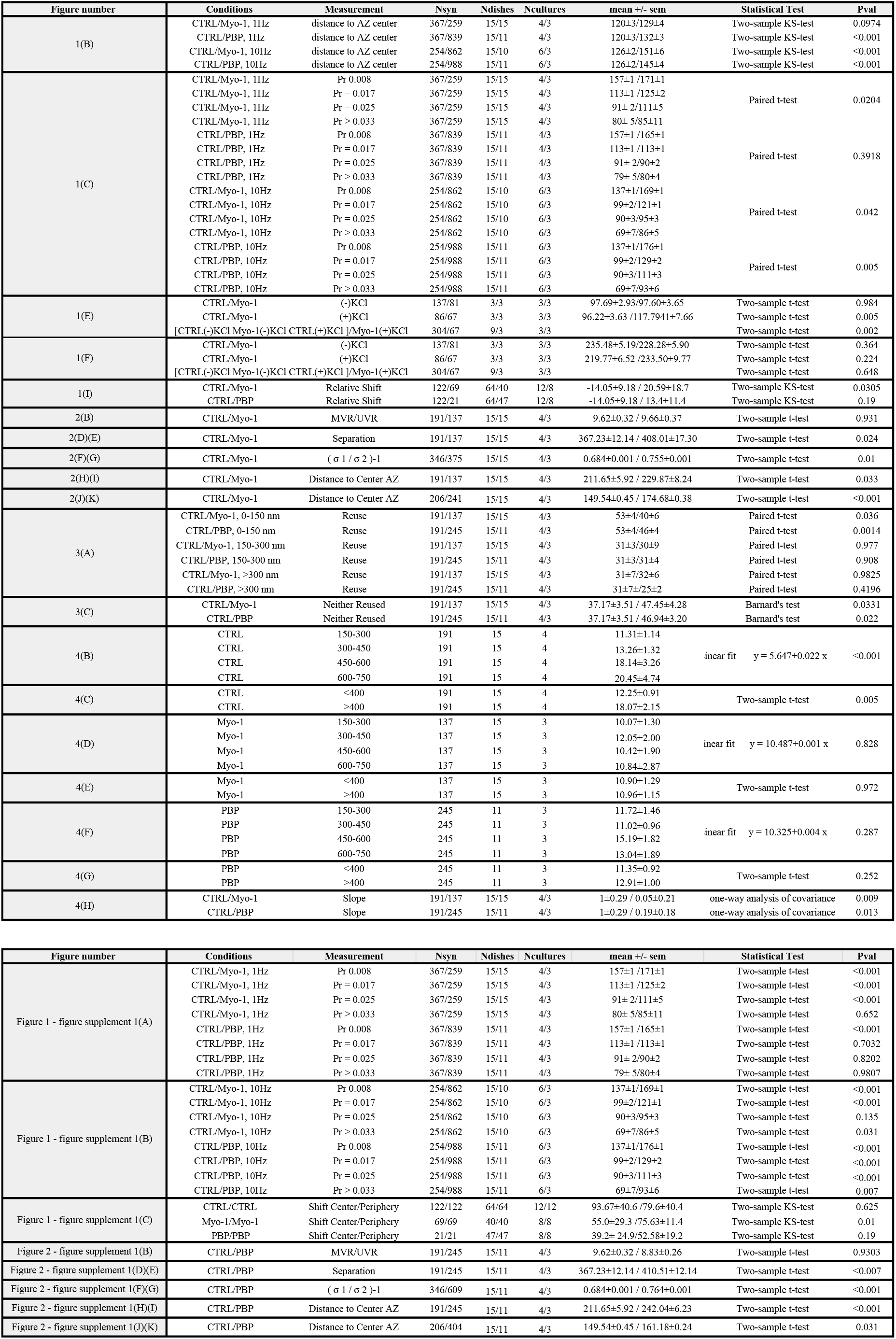
Table of all data values and statistical analyses. Data table columns are formatted as (i) corresponding figure location; (ii) conditions being statistically compared and separated by ‘/’; (iii) measurement (iv) number of samples (synapses, dishes, cultures) used for each test; (v) mean values and errors for each condition separated by ‘/’ and corresponding to conditions in column (i); (vi) statistical test used for comparison; (vii) P-value resulting from the statistical comparison.

To understand how this spatial shift arises, we examined changes in release site utilization upon myosin V inhibition. Individual release sites within each bouton were defined using hierarchical clustering algorithms with a cluster diameter of 50 nm (**Figure 1A)** as we described previously (Maschi and Klyachko, 2017). The observed spatial distribution of vesicle fusion events reflects a ~4-fold gradient of release site usage within the individual AZs, in which release sites with higher release probability are localized closer to the AZ center, while the sites that are used less frequently are localized more peripherally (**Figure 1C, Figure 1—figure supplement 1A)**. Most importantly, acute inhibition of myosin V resulted in a shift of release site utilization from the AZ center towards periphery at 1Hz and particularly at 10Hz stimulation **(Figure 1C; Figure 1—figure supplement 1A-B; Table 1)**, suggesting a role for myosin V in spatially controlling synaptic vesicle release.

To better understand the role of myosin V in spatial distribution of release, we analyzed the scanning electron microscopy (LaSEM) images of primary cultures of hippocampal neurons that were incubated (or not) with Myo-1 for 20 min and then acutely depolarized (or not) with 55 mM KCl for 10 min to induce vesicle release and recycling (Maschi et al., 2018). We examined both ‘docked’ vesicles (previously defined as vesicles with the center within 30 nm from the AZ), and ‘tethered’ vesicles (all vesicles with a center within 100 nm from the AZ) **(Figure 1D)**. Within these definitions, we found that myosin V inhibition selectively affected the spatial distribution of “docked” vesicles, causing a significant increase in the distance of docked vesicles from the AZ center upon KCl stimulation **(Figure 1E; Table 1).** This spatial shift in the localization of vesicles undergoing recycling and re-docking is in line with the spatial shift in the utilization of release sites caused by myosin V inhibition **(Figure 1C; Table 1)**. In contrast, no significant effect of Myosin V inhibition was observed in the absence of stimulation **(Figure 1E; Table 1)**, or within the “tethered” vesicle population in either condition **(Figure 1F; Table 1)**, suggesting the specific effects of myosin V inhibition on vesicle re-docking.

To further support these observations, we performed spatial analyses of the tracks of individual synaptic vesicles during recycling and docking in live hippocampal boutons, which we previously recorded in the presence (or not) of myosin V inhibitors (Maschi et al., 2018). Briefly, individual vesicles were labeled with a lipophilic FM-like dye SGC5 via compensatory endocytosis using a pair of stimuli at 100 ms. Single-vesicle tracking approach permitted us to follow the dynamics of individual vesicles with ~20 nm precision. We previously observed that vesicles undergo rounds of docking/undocking and accompanying transitions between the membrane pool and the inner synaptic pool. These transitions are evident as disappearance and reappearance events when vesicles are moving out-of and in-to the field of view near the AZ (**Figure 1G**) (Maschi et al., 2018). We thus quantified how myosin V inhibition affects the change in vesicle docking position by comparing its initial position before undocking/disappearance (ρ, **Figure 1G,H**) and its subsequent position upon reappearance/re-docking ( i.e. relative shift in docking location: Δ, **Figure 1G,H**). We observed that in control conditions, vesicles have a tendency to re-appear slightly closer to the synapse center, resulting in a net negative re-appearance shift in location relative to their original docking location (**Δ**= −14 ± 9 nm, see Methods for definition) **(Figure 1I; Table 1**), which is in line with the notion that more central release sites are preferentially utilized under basal conditions. In contrast, acute inhibition of myosin V with Myo-1 lead to a significant shift in relative vesicle re-docking position toward the synapse periphery upon re-appearance, resulting in a net positive re-appearance shift (**Δ**= +21 ± 18 nm; P = 0.03, two-tailed KS-test as compared to control condition) **(Figure 1I; Figure 1—figure supplement 1C; Table 1**). PBP treatment also showed a tendency of vesicle re-docking to occur more peripherally, but this effect was not statistically significant **(Figure 1I; Figure 1—figure supplement 1C; Table 1**). These differences could reflect the fact that the two agents have different mechanisms of action (Bond et al., 2013) and thus different effects on vesicle mobility: Myo-1 inhibits ADP release from actomyosin complex thus arresting myosin V on actin, while PBP reduces myosin-actin coupling by inhibiting ATP binding and hydrolysis; thus the two agents differentially affect the initial vesicle mobility state. Notably the vesicle tracking measurements are also not equivalent or directly comparable to the EM measurements or the vesicle release measurements above, because in our measurements vesicle displacement can only be defined relative to the 2D projection of the synapse center (as approximated by the geometric center of the total labeled recycling vesicle population, see Methods), but not the actual AZ center. Nevertheless, the spatial shift in vesicle re-docking position towards synapse periphery upon myosin V inhibition supports the other two experimental observations that myosin V modulates the spatial location of vesicle docking.

### Spatial organization of MVR events is myosin-V dependent

Analyses presented above have thus far examined the effects of myosin V inhibition on spatial properties of UVR. Additionally, MVR is also a prominent form of synchronous release in central synapses. We previously showed that the spatiotemporal organization of MVR events is determined by the gradient of release probability across the AZ (Maschi and Klyachko, 2020). Since myosin V supports refilling of individual release sites, we hypothesized that it could also regulate the spatial organization of MVR. To approach this question, we detected and analyzed individual MVR events in the same dataset that we used for analyses of UVR events above, as we described previously (Maschi and Klyachko, 2020). Briefly, in our recordings the vast majority of MVR events are evident as a pair of fusion events evoked by a single AP. Depending on the distance between the two vesicle fusion events comprising an MVR, such events fall in two subcategories. First subcategory contains well-separated MVR events that have sufficient spatial separation to allow each event in the pair to be individually localized (Resolved events, **Figure 2A)**. The second subcategory contain strongly overlapping, sub-diffraction distance MVR events that could not be resolved directly (Unresolved events), which required an alternative analysis approach comprising two separate steps. First, MVR event detection was achieved based on their amplitude (with a threshold set at two standard deviations above the mean quantal event amplitude determined individually for each bouton). Second, the identified MVR events were analyzed on the basis of asymmetry considerations, using an asymmetric Gaussian model fit to determine the width (sigma) of the Gaussian fit in the maximal (longitudinal, δ1) direction and the minimal (transverse, δ2) direction (**Figure 2F**, insert). The ratio δ1/δ2–1 (asymmetry score) represents asymmetry of the double-event image, which correlates with the distance between the two sub-diffraction events forming the image (DeCenzo et al., 2010). We have previously shown that the two subcategories have the same spatiotemporal features and represent the same biological phenomenon of MVR (Maschi and Klyachko, 2020).

**Figure 2:**
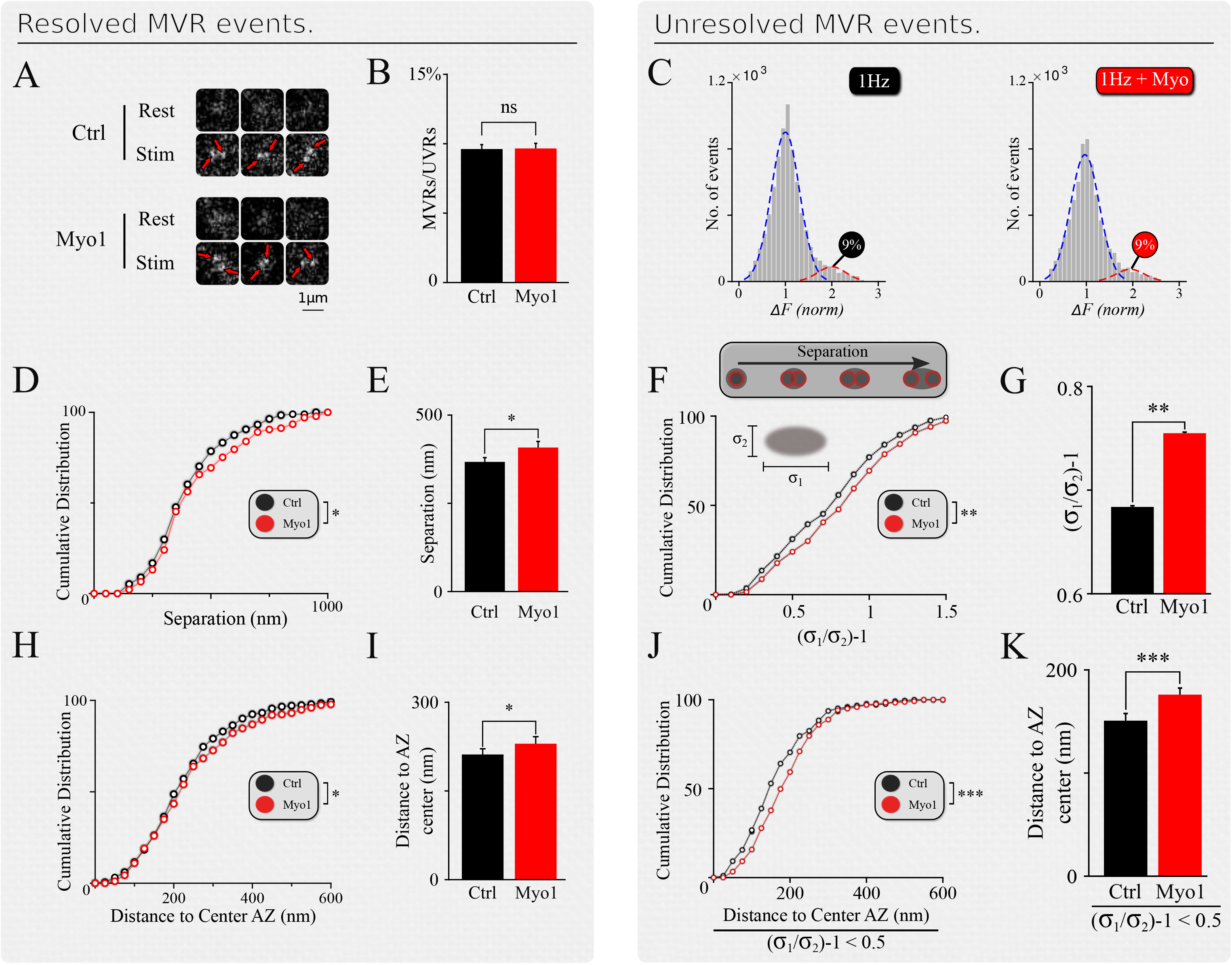
Spatial organization of MVR events is myosin-V dependent. (**A**) Examples of resolved MVR events in different boutons in control conditions (top) and cultures treated with Myo1 (bottom) for 20 min. Scale bar 1μm. (**B, C**) Inhibition of myosin V with Myo-1 does not affect the ratio between detected MVR and UVR events for resolved (**B**) and unresolved (**C**) MVR events. For unresolved MVR events, ratio of UVR and MVR was calculated based on a multi-Gaussian fit (**C**). **(D, E)** Effects of myosin V inhibition with Myo-1 on the distance between two fusion events comprising an MVR for resolved events. Cumulative plots (**D**) and mean values (**E**) are shown. **(F, G)** Same as (**D, E**) for unresolved MVR events. **(H, I)** Effects of myosin V inhibition with Myo-1 on the distance from MVR events to the AZ center for resolved events. Cumulative plots (**H**) and mean values (**I**) are shown. **(J, K)** Same as (**H, I**) for unresolved MVR events. Only a subpopulation of more symmetrical MVR events (asymmetry score<0.5) were included in this analysis, because these more symmetrical events could be well-approximated by a single symmetrical Gaussian fit, making this analysis comparable to that of the resolved MVR events. Two-sample t-test (all panels). *p<0.05, **p<0.01, ***p<0.001, ns - not significant.

Inhibition of myosin V did not strongly affect the UVR/MVR event ratio for either population of resolved or unresolved MVR events **(Myo-1: Figure 2B,C; PBP: Figure 2— figure supplement 1B,C; Table 1)**. However several spatial features of MVR were affected by myosin V inhibition. First, the separation distance between the two releases comprising an MVR event was significantly increased in the presence of Myo-1 or PBP, for both resolved and unresolved MVR events **(Myo-1: Figure 2D,E and 2F,G; PBP: Figure 2—figure supplement 1D,E and 1F,G; Table 1)**. Second, both resolved and unresolved MVR events occurred further away from the AZ center when myosin V was inhibited **(Myo-1: Figure 2H,I and 2J,K; PBP: Figure 2—figure supplement 1H,I and 1J,K; Table 1)**. These results are consistent with the above notion that myosin V inhibition causes a shift in utilization of release sites away from the AZ center.

To confirm and further explore the role of myosin V in the spatial aspects of release site utilization we analyzed the reuse of the release sites engaged in MVR. We observed that central release sites engaged in MVR events show a significant reduction of reuse upon myosin V inhibition, while the more peripheral release sites were not strongly affected **(Figure 3A; Table 1)**. This observation thus provides a mechanistic basis for the increased distance from the MVR events to the AZ center and the correspondingly increased spatial separation within the MVR event pair that we observed above **(Figure 2; Table 1)**.

**Figure 3:**
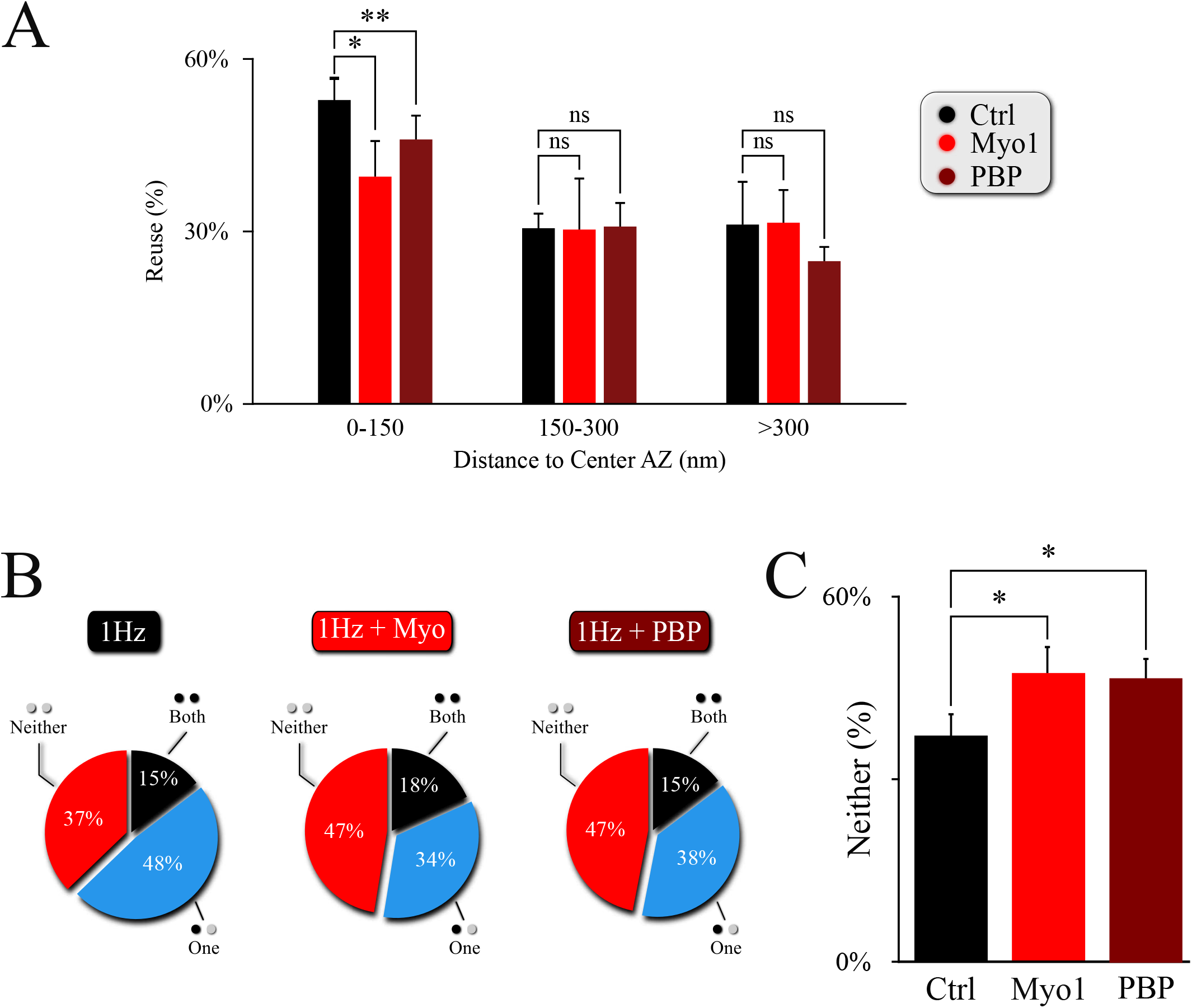
Myosin V regulates utilization of release sites engaged in MVR. (**A**) Effect of myosin V inhibition with Myo-1 (red) or PBP (brown) on reuse of release sites engaged in MVR events. Reuse was quantified as the percentage of release sites engaged in MVR that are reused at least once during the 120sec observation period by either other MVR or UVR events. The reuse probability is highly dependent on the distance to AZ center; to account for this variability, we use a paired t-test with data binning at 50 nm. (**B, C**) Effect of myosin V inhibition with Myo-1 (red) or PBP (brown) on the spatial overlap of MVR and UVR events determined by the proximity analysis. Percentages of MVR events in which none (red), one (blue) or both (black) events in the pair occurred within ±25nm of at least one UVR event (i.e. at the same at release site) during the observation period (**B**), and quantification of the percentage of no overlap of MVR and UVR events (**C**). Two-sample t-test (A) or Barnard’s test (C). *p<0.05, ns - not significant.

This observation also provides a testable prediction. We previously found that release sites closer to the AZ center are more likely to harbor UVR as well as MVR events during observation period (representing spatial “overlap” of UVR and MVR events). Thus reduced utilization of the central release sites upon myosin V inhibition predicts that the spatial overlap of UVR and MVR events at the same release sites is also reduced. To test this prediction, we analyzed the probability that the same release site is engaged in UVR and MVR during our observation time. As predicted, the overlap of MVR and UVR events at the same release sites was significantly reduced in the presence of Myo-1 or PBP (**Figure 3B,C, Table 1**).

Therefore, by reducing the preferential utilization of central release sites during MVR, inhibition of Myosin V not only results in increased distance from MVR events to the AZ center and increased spatial separation within individual MVR events, but it also reduces spatial overlap of MVR with the UVR events.

### Inhibition of myosin V reduces temporal separation within MVR events

The pairs of release events comprising MVR are often not perfectly synchronized with each other, but exhibit a slight temporal separation on the order of 1-5 milliseconds (Auger et al., 1998; Auger and Marty, 2000; Crowley et al., 2007; Malagon et al., 2016; Maschi and Klyachko, 2020; Rudolph et al., 2011). We recently showed that this temporal separation arises because the first event in the MVR pair occurs closer to the AZ center, while the second event in the pair occurs more peripherally with a slight delay (Maschi and Klyachko, 2020). The extent of this temporal separation depends on the difference in radial distance of the two events comprising MVR from the AZ center and correlates with the distance between the two events **(Figure 4B; Table 1)**. Because the spatial localization of MVR events is altered by myosin V inhibition, we examined how the temporal separation is affected. To estimate the temporal separation within the MVR events we measured the amplitude differences between the two events in the same frame, which is an established approach to quantify the temporal separation (Maschi and Klyachko, 2020) **(Figure 4A)**. Here we found that in the presence of Myo-1 (**Figure 4D,E)** or PBP **(Figure 4F,G)**, the amplitude differences within the individual MVR events were no longer dependent on their relative distance (as compared to control, **Figure 4B, C**, and quantified in **Figure 4H; Table 1)**. We note that a component of the amplitude differences likely arises from an uncertainty in determining the fusion event amplitude; which we previously estimated to be ~10%. Thus, the amplitude differences remaining in our measurements in the presence of Myo-1 or PBP could be, to a large extent, accounted for by the intrinsic uncertainty in our measurements. These results suggest that inhibition of myosin V reduces the temporal separation within the MVR events. Thus myosin V regulates both spatial and temporal organization of MVR events as well as UVR.

**Figure 4:**
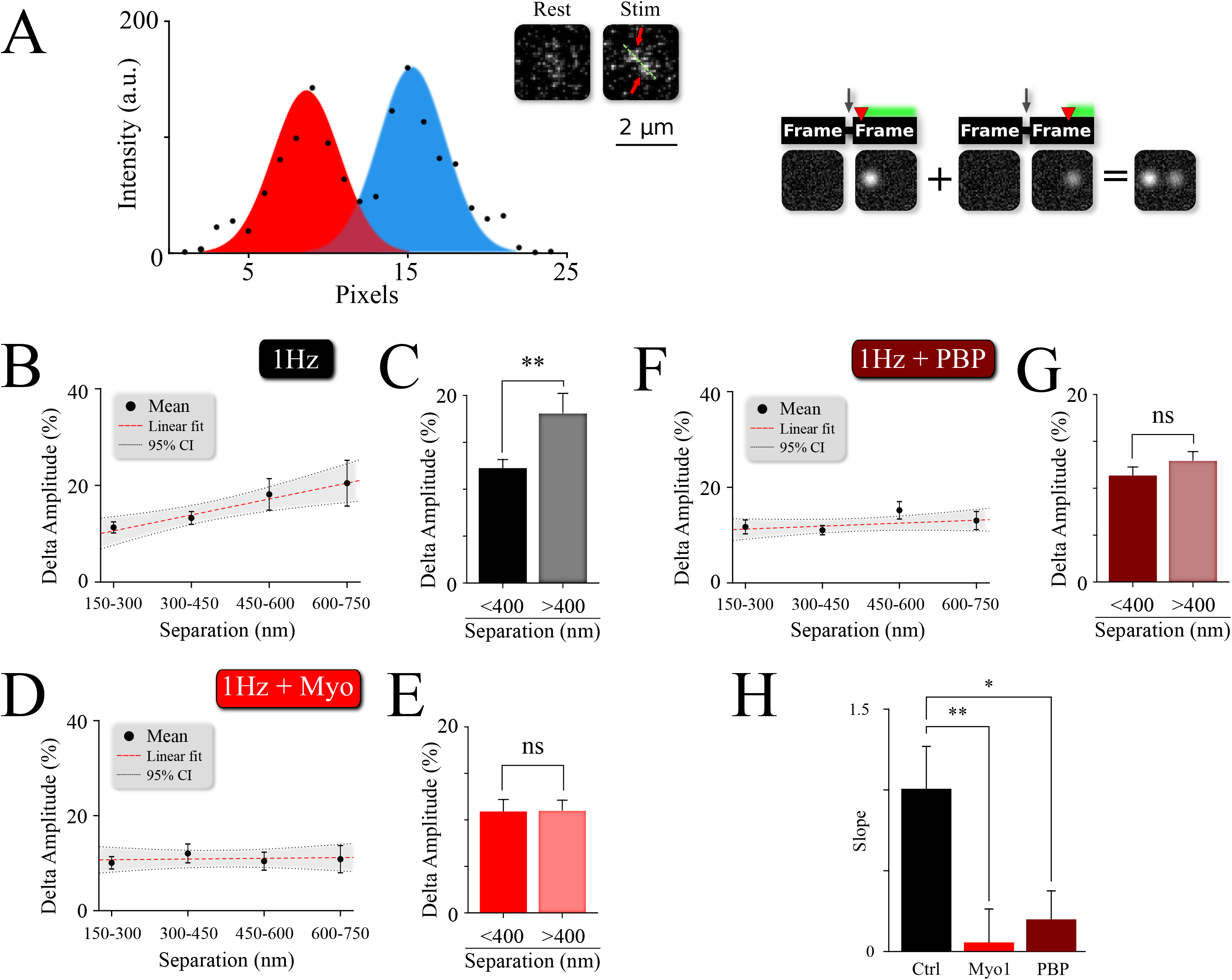
Myosin V regulates temporal separation within MVR events. **A**) *Left panel:* Sample image and intensity profile of an MVR event with noticeable difference in intra-event amplitude. *Right panel:* A cartoon diagram shows the relationship between time delay (red arrow) of the second fusion after an action potential and the resulting amplitude difference within an MVR event pair recorded together in the same frame. (**B, C**) Amplitude difference of the two events comprising MVR as a function of intra-event distance. Linear fit (**B**) and t-test of pooled data (**C**) are shown. (**D, E)** Effect of myosin V inhibition with Myo-1 on the amplitude difference of the two events comprising MVR as a function of intra-event distance. Linear fit (**D**) and t-test of pooled data (**E**) are shown. **(F, G**) Same as (**D,E**) for the effect of myosin V inhibition with PBP. (**H**) Quantification of the effects of Myo-1 or PBP in panels (**D**) and (**F**) assessed by comparing the slopes of the correlations in (**B**), (**D**) and (**F**). Two-sample t-test (C, E, G, H). *p<0.05, **p<0.001. ns – not significant.

## DISCUSSION

Docking of synaptic vesicles at the release sites within the AZ is an essential mechanism controlling strength and timing of synaptic transmission. We previously showed that vesicle-associated molecular motor myosin V is a key regulator of release site refilling during synaptic activity by controlling vesicle anchoring and retention at the release sites. Here we extend these studies to demonstrate that myosin V also regulates the spatial organization of vesicle docking across the AZ during two main forms of synchronous release, the UVR and MVR. This is supported by three key observations: (i) Acute inhibition of myosin V shifts location of vesicle docking away from the AZ center towards periphery. Consequently the utilization of release sites during UVR also shifts away from the AZ center when myosin V is inhibited; (ii) Inhibition of myosin V reduces utilization of central release sites by MVR events. Consequently MVR events occur further away from the AZ center and have a larger separation distance within the event pair; (iii) Inhibition of myosin V reduces the temporal separation within the MVR events. Thus by regulating spatio-temporal organization of UVR and MVR events across the AZ, myosin V actions represent a mechanism that fine-tunes neurotransmitter release.

### Myosin V role in the spatiotemporal regulation of UVR and MVR

The spatial and temporal utilization of release sites during both UVR and MVR follows complex patterns that are determined by the gradient of release probability (Pr) across the AZ. Yet such apparent complexity often arises from simpler underlying principles thus posing a central question: given the function of Myosin V in vesicle anchoring/docking at release sites, could the observed effects of myosin V inhibition on release site utilization be explained simply by changes in the gradient of release site Pr? To approach this question, we created a basic model representation of an AZ with 12 discrete release sites arranged to form a center-to-periphery gradient of release probability (Pr) (**Schematic 1A**). Because the number of release sites per AZ vary widely across synapse population (in the range of 2-18 (Maschi and Klyachko, 2017; Sakamoto et al., 2018; Tang et al., 2016)), the model was formulated not to depend on the precise number of release sites, but rather on the gradient of release site Pr (central/peripheral) across the AZ. First, the model shows that reducing the center-to-periphery gradient of Pr across the AZ results in increased distance of UVR events to the AZ center (**Schematic 1B**), which is what we observed experimentally as a result of myosin V inhibition. Likewise, for the MVR events, the model shows that reducing the Pr gradient also leads to increased spatial separation of the two fusion events comprising an MVR (**Schematic 1C**), which we also observed following myosin V inhibition. Thus the simplest working model that accounts for the observed spatial effects of myosin V inhibition is that by shifting utilization of release sites from more central to more peripheral, myosin V inhibition acts by reducing the Pr gradient effectively spreading the release to a larger area of the AZ.

**Schematic 1:**
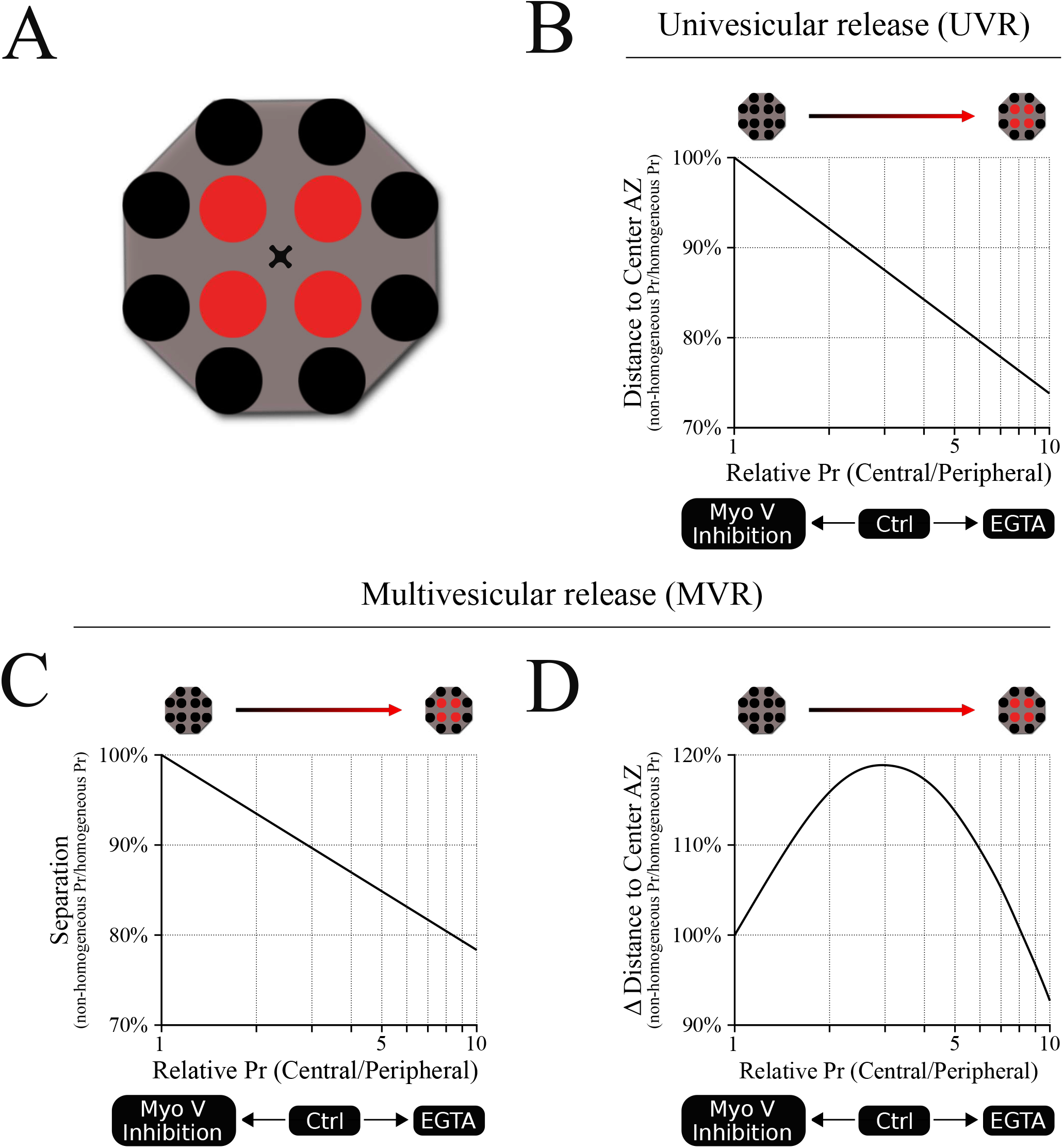
A model linking release site Pr gradient and spatiotemporal features of UVR and MVR events. (**A**) A cartoon representation of the model illustrating spatial distribution of 12 release sites within a single AZ. We used a Monte Carlo simulation to model the probability that a release event occurs in a given release site based on the release probability of the individual release sites. For this model, a shared probability was assigned for the four central release sites **(red)** and a different but also shared probability was assigned for the 8 peripheral release sites **(black)**. In other words, the model could be represented by two concentric donuts with two different Pr values. Ten different central/peripheral Pr ratios (i.e. Pr gradients) were used (from 1 to 10); for each Pr ratio we run 1 million simulations, resulting in the outcome of 10 million points for each plot shown. The results were normalized to the values obtained at the Pr ratio = 1 (homogeneous distribution of Pr across the AZ). (**B**) Release events occur at shorter distances from the AZ center as the central/peripheral Pr ratio increases; in other words, the utilization of more central release sites increases as the Pr gradient increases. The opposite effect occurs when Pr ratio decreases, leading to larger distances from release events to the AZ center, representing an increased utilization of more peripheral release sites. (**C**) Spatial separation within the pair of release events comprising an MVR decreases as the central/peripheral Pr gradient increases. The opposite occurs when Pr gradient decreases, leading to increased spatial separation within the MVR events, as we observed when myosin V was inhibited. (**D**) Temporal separation between the two release events comprising an MVR depends on the difference in their radial distances to AZ center. This parameter has a bell-shape dependence on the central/peripheral Pr ratio. Either increase or decrease of the Pr ratio from the optimal value around 3-4 leads to smaller differences in the distances to the AZ center for the two events comprising an MVR. This predicts a reduced temporal separation of the two events comprising an MVR with either increase or decrease in the Pr gradient, as observed experimentally with EGTA or inhibition of myosin V, respectively.

The conceptual relationship between the steepness of the Pr gradient and spatial localization of release events also holds under conditions when Pr gradient becomes steeper than normal. For example, we previously observed that buffering intraterminal calcium with EGTA increased utilization of central release sites (thus making the center-to-periphery Pr gradient steeper). EGTA also caused a shift in spatial localization of UVR events towards the AZ center and reduced spatial separation within the MVR events (Maschi and Klyachko, 2017,2020), both of which are recapitulated by the model (**Schematic 1B,C**).

Moreover, this framework also recapitulates the more complex relationship between the Pr gradient and the temporal features of MVR. Interestingly, both inhibition of myosin V and buffering intraterminal calcium with EGTA have the same effect of decreasing the temporal separation within MVR events, while having opposing effects on the Pr gradient. While appear paradoxical on the first glance, these results are also conceptually explained by our model. Specifically, our previous observations suggested that temporal separation within MVR events results from the first event occurring closer to the AZ center, while the second event in the pair occurring with a short 1-5ms delay at a more peripheral site. This temporal separation thus depends on the difference in radial distance to AZ center of two fusion events comprising an MVR. Our model shows that this parameter has a bell-shape curve (**Schematic 1D**), reaching a maximum at ~3-4-fold gradient of Pr, which is similar to experimentally observed value in control conditions. Therefore either inhibition of myosin V or calcium buffering with EGTA, while having opposite effects on the steepness of the Pr gradient, both drive it away from the optimal value, resulting in reduced temporal separation. This temporal control, in combination with regulating spatial separation within the MVR events, may allow myosin V to fine-tune the quantal size by adjusting the duration of neurotransmitter release during MVR while engaging spatially distinct subsets of postsynaptic receptors.

### Myosin V and the gradient of release site properties

Our results do not necessarily imply that myosin V selectively serves as a docking factor only for the central release sites; the effect of myosin V inhibition could simply be more apparent for the central release sites because they are used much more frequently under normal conditions, while the limited duration of observation masks the effect on peripheral release sites which are used much less frequently. Thus we speculate that additional or alternative mechanisms may exist that makes usage of central release sites more frequent. One possible mechanism suggested by our previous study is the presence of center-to-periphery gradient of calcium elevation following an action potential (Maschi and Klyachko, 2020). Such calcium gradient could in turn control myosin V-dependent vesicle retention at release sites thus creating a center-to-periphery gradient of release site Pr. Indeed, Myosin V function is calcium-dependent; calcium elevation drives transition of myosin V from a transporting motor to a tether and also regulates myosin V association with the SNARE proteins (Krementsov et al., 2004; Ohyama et al., 2001; Prekeris and Terrian, 1997; Watanabe et al., 2005). Thus the differences in spatial utilization of release sites could be driven by the gradient of calcium elevation in the synaptic bouton following an AP, which determines the strength or duration of myosin V association with a release site. While the mechanistic basis for the gradient of calcium rise across the AZ will require further investigation, a number of possible mechanisms have been suggested in previous studies. A higher calcium elevation in the AZ center can simply result from larger density of release sites (assuming each is associated with a calcium channel) at the AZ center vs periphery. Differential calcium channel mobility in the center vs periphery of the AZ (Schneider et al., 2015) could also contribute to different stability of channel association with the release sites or its coupling with the vesicle (Eggermann et al., 2011; Miki et al., 2017). Alternatively, or additionally, a gradient of release site properties could arise from other, calcium/myosin V- independent mechanisms. For instance clusters of presynaptic proteins that are believed to represent the structural correlates of release sites exhibit a large degree of heterogeneity in size and composition across the AZ (Glebov et al., 2017; Schneider et al., 2015; Tang et al., 2016) presumably due to differential enrichment and mobility of many critical components, such as Bassoon, RIM, Munc13, Munc18, and Syntaxin-1 (Bademosi et al., 2017; Glebov et al., 2017; Schneider et al., 2015; Smyth et al., 2013; Tang et al., 2016; Weyhersmuller et al., 2011). Clusters of several of these critical proteins are detected predominately near the AZ center (Tang et al., 2016), suggesting that more peripheral clusters are smaller and below the detection limit.

In summary, by modulating the landscape of release probability across the AZ, myosin V fine-tunes the spatio-temporal dynamics of neurotransmitter release during both UVR and MVR events to dynamically shape synaptic transmission.

## EXPERIMENTAL METHODS

### Neuronal Cell Cultures

Neuronal cultures were produced from the hippocampus of E16-17 rat pups of mixed gender as previously described (Maschi et al., 2018; Peng et al., 2012). Hippocampi were dissected from E16-17 pups, dissociated by papain digestion, and plated on coated glass coverslips containing an astrocyte monolayer. Neurons were cultured in Neurobasal media supplemented with B27. All animal procedures conformed to the guidelines approved by the Washington University Animal Studies Committee.

### Lentiviral Infection

VGlut1-pHluorin was generously provided by Drs. Robert Edward and Susan Voglmaier (UCSF) (Voglmaier et al., 2006). Lentiviral vectors were generated by the Viral Vectors Core at Washington University. Hippocampal neuronal cultures were infected at DIV3.

### Fluorescence Microscopy

#### Neurotransmitter Release Measurements

All experiments were conducted at 37°C within a whole-microscope incubator (In Vivo Scientific) at DIV16–19 as described previously (Maschi et al., 2018). Neurons were perfused with bath solution (125 mM NaCl, 2.5 mM KCl, 2 mM CaCl2, 1 mM MgCl2, 10 mM HEPES, 15 mM Glucose, 50 mM DL-AP5, 10 mM CNQX, pH adjusted to pH 7.4). Fluorescence was excited with a Lambda XL lamp (Sutter Instrument) through a 100x 1.45 NA oil-immersion objective and captured with an EMCCD camera (Hamamatsu) or cooled sCMOS camera (Hamamatsu). Focal plane was continuously monitored, and focal drift was automatically adjusted with 10 nm accuracy by an automated feedback focus control system (Ludl Electronics). Field stimulation was performed by using a pair of platinum electrodes and controlled by the software via Master-9 stimulus generator (A.M.P.I.). Images were acquired using two frames with an acquisition time of 40ms, one 45ms before stimulation and one coincidently (0ms delay) with stimulation.

#### Single-vesicle tracking

Sparse vesicle labeling and functional synapse localization were performed following our previously developed procedures (Maschi et al., 2018). The same bath solution as above was used for the dye loading and imaging, except 0.2 mM CaCl_2_, 1.0 mM MgCl_2_ were used to wash excess dye from the sample. 10 μM SGC5 (Biotium) were added to the bath solution for the dye loading step. Samples were imaged for 50-70 sec, at an exposure rate of 80 msec (with a total frame rate of 10Hz). Samples were stimulated for 10 sec at 20 Hz with a 10 sec delay after the first frame.

### Pharmacology

MyoVin-1 (Millipore), Pentabromopseudalin (PBP, Fisher Scientific) or EGTA-AM (Millipore) were diluted in DMSO (Sigma-Aldrich) and stored at −20°C. Samples were incubated in imaging solution with 30 μM Myo-1 for 5-10 min or 5 μM PBP for 5 min, or 250 μM EGTA-AM for 20 min before dye loading. The effective final DMSO concentration was <0.5%. Extended exposure to MyoVin-1 or PBP caused cell death, thus the bath solution during the experiment did not include Myo-1 or PBP. Our control measurements indicated that continuous presence of these blockers during the experiments did not have additional effects on vesicle motility beyond the effects of pre-incubation (data not shown).

### Large-Area Scanning Electron Microscopy (LaSEM)

Cultures were fixed in a solution containing 2.5% glutaraldehyde and 2% paraformaldehyde in 0.15 M cacodylate buffer with 2 mM CaCl2, pH 7.4 that had been warmed to 37°C for one hour. In experiments with KCl-induced depolarization, fixation was performed immediately following KCl application, and care was taken to complete the fixation procedure within a few seconds. Coverslips were rinsed in cacodylate buffer 3 times for 10 minutes each, and subjected to a secondary fixation for one hour in 2% osmium tetroxide/1.5% potassium ferrocyanide in cacodylate buffer for one hour, rinsed in ultrapure water 3 times for 10 minutes each, and stained in an aqueous solution of 1% thiocarbohydrazide for one hour. After this, the coverslips were once again stained in aqueous 2% osmium tetroxide for one hour, rinsed in ultrapure water 3 times for 10 minutes each, and stained overnight in 1% uranyl acetate at 4°C. The samples were then again washed in ultrapure water 3 times for 10 minutes each and en bloc stained for 30 minutes with 20 mM lead aspartate at 60°C. After staining was complete, coverslips were briefly washed in ultrapure water, dehydrated in a graded acetone series (50%, 70%, 90%, 100% x2) for 10 minutes in each step, and infiltrated with microwave assistance (Pelco BioWave Pro, Redding, CA) into Durcupan resin. Samples were flat embedded in a polypropylene petri dish and cured in an oven at 60°C for 48 hours. Post resin curing, the coverslips were exposed with a razor blade and etched off with concentrated hydrofluoric acid. Small pieces of the resin containing the cells was then cut out by saw and mounted onto blank resin stubs before 70 nm thick sections were cut in the cell culture growing plane and placed onto a silicon wafer chips. These chips were then adhered to SEM pins with carbon adhesive tabs and large areas (~ 330 × 330 μm) were then imaged at high resolution in a FE-SEM (Zeiss Merlin, Oberkochen, Germany) using the ATLAS (Fibics, Ottowa, Canada) scan engine to tile large regions of interest. High-resolution tiles were captured at 16,384 × 16,384 pixels at 5 nm/pixel with a 5 μs dwell time and line average of 2. The SEM was operated at 8 KeV and 900 pA using the solid-state backscatter detector. Tiles were aligned and export using ATLAS 5.

### Image and Data Analysis

#### Localization of UVR events

The fusion event localization at subpixel resolution was performed using MATLAB code based on the uTrack software package (Aguet et al., 2013; Jaqaman, 2008). Release sites were defined using hierarchical clustering performed in MATLAB as we described previously (Maschi et al., 2018; Maschi and Klyachko, 2017, 2020). We previously found that the observed clusters do not arise from random distribution of release events, but rather represent a set of defined and repeatedly reused release sites within the AZs (Maschi and Klyachko, 2017).

#### Localization of MVR events

Localization of resolved MVR events was performed using a mixture-model multi-Gaussian fit using in-built functions in uTrack (Aguet et al., 2013; Jaqaman, 2008) as we described previously (Maschi and Klyachko, 2020).

Unresolved MVR events were identified based on the event amplitude. The single event amplitude and its variability were determined for each bouton individually. Photobleaching was accounted for by fitting the event intensity changes over time. The threshold for MVR event detection was set at two standard deviations above the mean single event amplitude determined individually for each bouton. Localization of unresolved MVR events was determined using an asymmetrical Gaussian model fit based on the minimization of the residuals as described in (Maschi and Klyachko, 2020).

#### Release site reuse and release probability

Release probability of individual release sites was calculated based on the number of release events detected per release site and divided by the duration of the observation period. For MVR events, reuse was defined more broadly as the probability that the release site engaged in MVR is reused at least once during the 120sec observation period by either other MVR or UVR events.

#### Event proximity analysis

To determine probability of spatial overlap of MVR and UVR events at the same release sites during the observation period, a proximity analysis was performed in which overlap was defined as having at least one UVR event occurring within 25nm of an MVR event during observation period.

#### EM analyses

Synapse identification and vesicle localization analysis were performed as described in (Maschi et al., 2018). Distances to the AZ center were measured from the projection of the vesicle position on the AZ plane. “Docked” vesicles were defined as those with the distance from the membrane to the vesicle center less than 30 nm and “tethered” vesicle as those with the distance less than 100 nm.

#### Single-Vesicle Tracking

Individual vesicle track positions (x,y) were obtained using the MATLAB code based on uTrack software (Jaqaman, 2008) following our previously developed procedures (Forte et al., 2017; Gramlich and Klyachko, 2017; Maschi et al., 2018). Quantification of vesicle motion was performed using the three-frame moving average of vesicle position to mitigate the effects of noise. Vesicle tracks were converted from two-dimensional (x,y) spatial locations in the imaging plane to a one-dimensional radial distance from the synapse center (x_s_, y_s_), s = sqrt((x − x_s_)^2^ + (y− y_s_))^2^. Synapse center was defined as a center of mass of the synapse image obtained following labeling the entire vesicle population with a strong stimulus of 400APs at 20Hz. When more than one disappearing and/or re-appearing tracks were observed in a given synapse, all tracks associated with the same bouton were grouped together to determine the criterion for analysis described below.

Vesicles were accepted for the analysis based on the following conditions: [i] a vesicle must be localized within 600 nm of a synapse center within the first 20 frames and must be observed for at least 50 frames before disappearing; [ii] the vesicle must have a disappearance and a reappearance events as we defined previously (Maschi et al., 2018) and be observed for at least 20 frames afterwards; [iii] if multiple re-appearance events occur for the same vesicle, each event is counted as a new re-appearance with the same requirements. Synapses where more than one vesicle was observed were excluded from this analysis. Relative shift in vesicle location upon disappearance and re-docking was quantified as the difference in radial distances of vesicle re-appearance and disappearance positions. Average vesicle position before disappearance was quantified for the first five seconds of the track (ρ). Average position for the re-appeared vesicle was quantified for the entire time the track re-appeared (t >2 sec). All vesicle shifts for each condition (Ctrl, Myo-1, PBP) were pooled and binned into 25 nm bin-size distributions centered around 0 nm. Each side of the distribution (representing a shift towards or away from the synapse center) was fit separately to an exponential decay and the overall shift was determined as the difference in the fit time courses.

#### Vesicle Disappearance and Appearance Oversampling Correction

Vesicle disappearance and appearance distributions were sampled at a rate of 10 frames per second. However, the typical disappearance rate was on the order of 1 vesicle per second (1 vesicle per 10 frames) resulting in significant oversampling. Thus, we averaged the oversampled distributions with a five-frame moving average and plotted every fifth data point. Further, we performed statistical analysis on the averaged data to prevent over-sampling bias of the statistics.

### Statistical Analyses

Statistical analyses were performed in Matlab. Statistical significance was determined using two tailed Student’s t-test, Kolmogorov-Smirnov (K-S) test, or a Barnard’s test where appropriate. Data is reported as mean ± SEM; or ±95% confidence interval; or ± residual sum of squares from fits to distributions, as indicated in the text, figure legends and **Table 1**. p < 0.05 was considered statistically significant. The number of experiments reported reflects the number of different cell cultures tested and is provided in **Table 1**. Statistical tests used to measure significance are indicated in each figure legend along with the corresponding significance level (p value). Analysis of the samples was not blinded to condition. Randomization and sample size determination strategies are not applicable to this study and were not performed.

## ACKNOWLEDGMENTS

This work was supported in part by grants to VAK from NINDS (R35). We acknowledge the assistance of Matthew Joens and Dr. James Fitzpatrick at the Washington University Center for Cellular Imaging in EM studies

## SUPPLEMENTARY FIGURE LEGENDS

**Figure 1-figure supplement 1:**
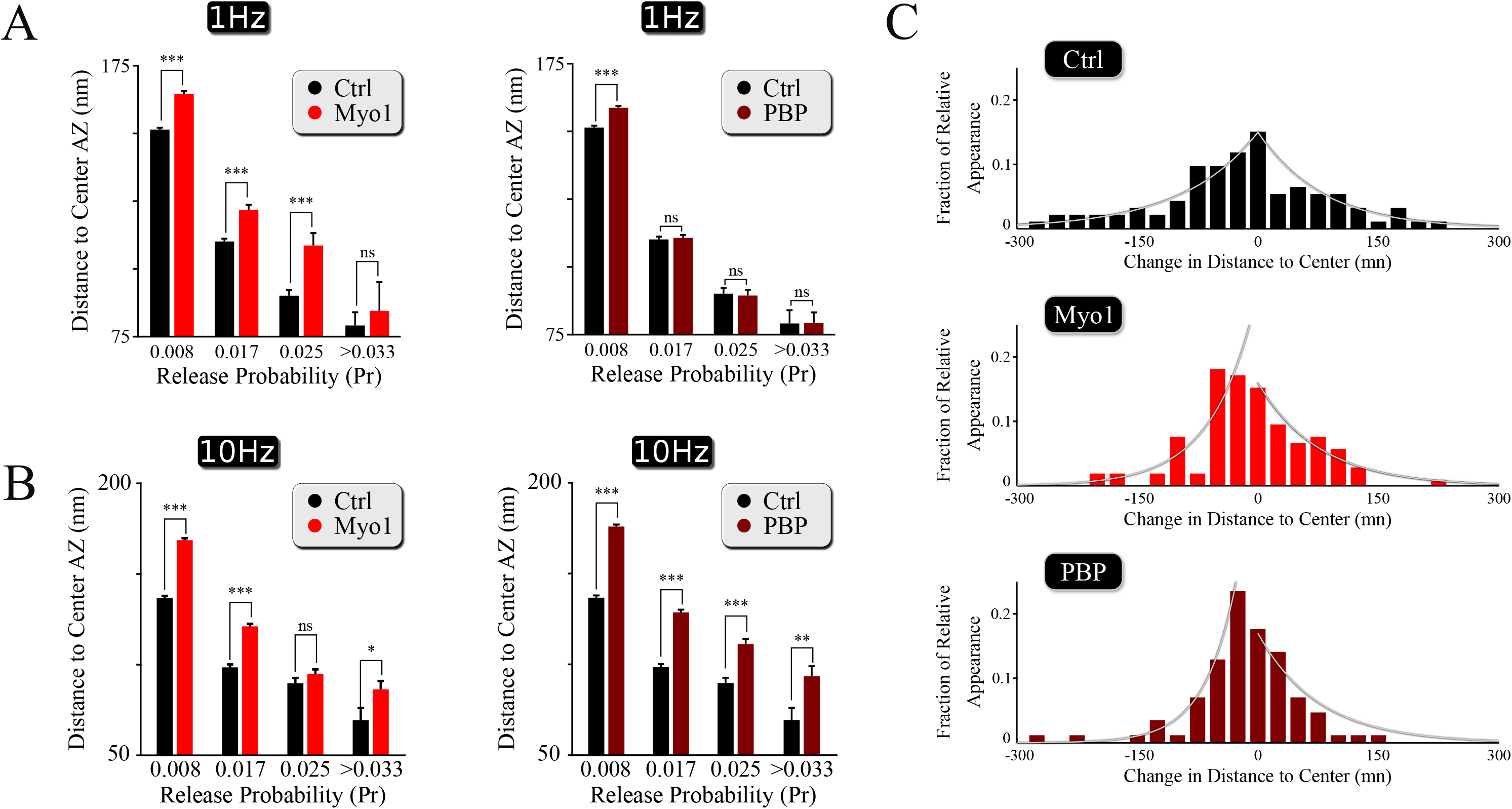
Spatial effects of myosin V inhibition on release site utilization and vesicle re-docking. (**A,B**) Effects of myosin V inhibition with Myo-1 (left) or PBP (right) on the average distance to the AZ center for individual release sites in measurements at 1Hz (top) or 10Hz (bottom), binned on the basis of their release probability. (**C**) Histograms of the shift in the distance to synapse center for vesicles undergoing a disappearance and a reappearance event in Control, Myo-1 and PBP. Locations of vesicle re-appearance were separated as toward synapse center versus toward periphery relative to the vesicle initial location (defined as a point of 0 shift) and each side of the histograms were fitted to a single exponential, the difference of which was used to determine the relative shift. Two-sample t-test (A, B). *p<0.05, **p<0.01, ns - not significant.

**Figure 2-figure supplement 1.**
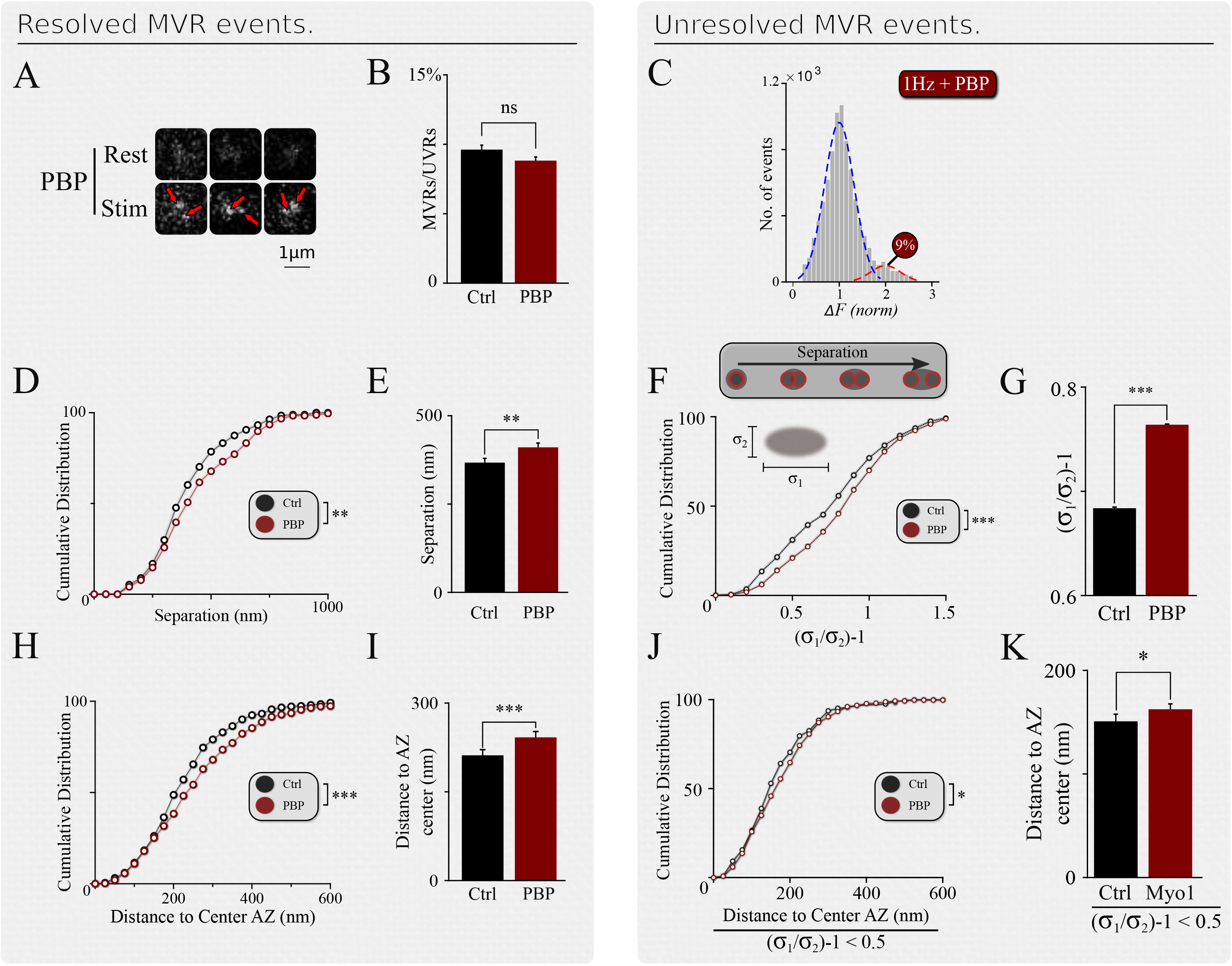
Effects of myosin V inhibition with PBP on the spatial organization of MVR events. (**A**) Examples of resolved MVR events in cultures treated with PBP. Scale bar 1μm. (**B, C**) Inhibition of myosin V with PBP does not affect the ratio between MVR and UVR events for resolved (**B**) and unresolved MVR (**C**) events. For unresolved MVR events, ratio of UVR and MVR was calculated based on a multi-Gaussian fit (**C**). **(D, E)** Effects of myosin V inhibition with PBP on the distance between two fusion events comprising an MVR for resolved events. Cumulative plots (**D**) and mean values (**E**) are shown. **(F, G)** Same as (**D, E**) for unresolved MVR events. **(H, I)** Effects of myosin V inhibition with PBP on the distance from MVR events to the AZ center for resolved events. Cumulative plots (**H**) and mean values (**I**) are shown. **(J, K)** Same as (**H, I**) for unresolved MVR events. Only a subpopulation of more symmetrical MVR events (asymmetry score<0.5) were included in this analysis, because these more symmetrical events could be well-approximated by a single symmetrical Gaussian fit, making this analysis comparable to that of the resolved MVR events. Two-sample t-test (all panels). *p<0.05, **p<0.01, ***p<0.001, ns - not significant.

---

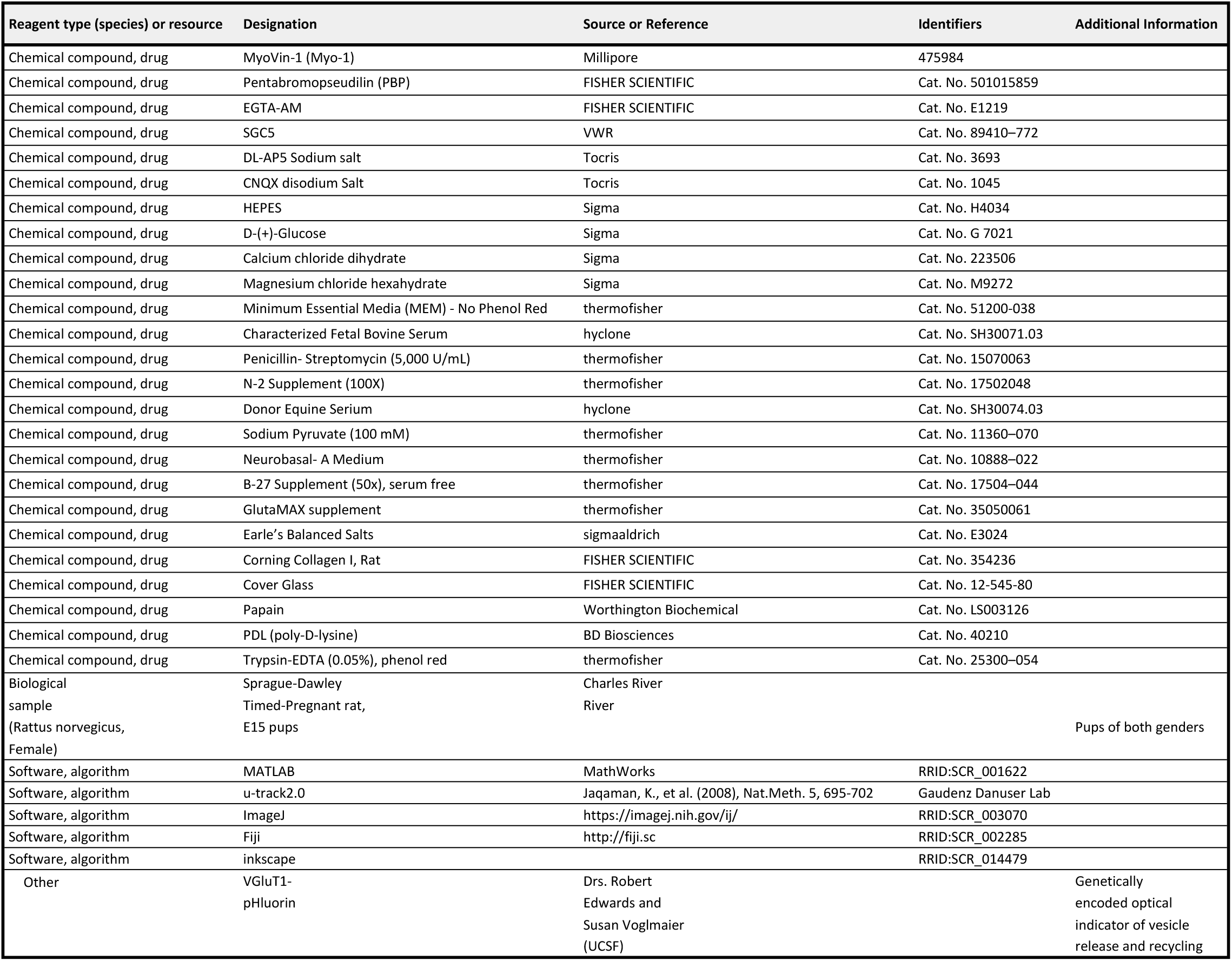
Key resources table.

Key resources table.
| Reagent type (species) or resource | Designation | Source or Reference | Identifiers | Additional Information |
| --- | --- | --- | --- | --- |
| Chemical compound, drug | MyoVin-1 (Myo-1) | Millipore | 475984 |  |
| Chemical compound, drug | Pentabromopseudilin (PBP) | FISHER SCIENTIFIC | Cat. No. 501015859 |  |
| Chemical compound, drug | EGTA-AM | FISHER SCIENTIFIC | Cat. No. E1219 |  |
| Chemical compound, drug | SGC5 | VWR | Cat. No. 89410–772 |  |
| Chemical compound, drug | DL-AP5 Sodium salt | Tocris | Cat. No. 3693 |  |
| Chemical compound, drug | CNQX disodium Salt | Tocris | Cat. No. 1045 |  |
| Chemical compound, drug | HEPES | Sigma | Cat. No. H4034 |  |
| Chemical compound, drug | D-(+)-Glucose | Sigma | Cat. No. G 7021 |  |
| Chemical compound, drug | Calcium chloride dihydrate | Sigma | Cat. No. 223506 |  |
| Chemical compound, drug | Magnesium chloride hexahydrate | Sigma | Cat. No. M9272 |  |
| Chemical compound, drug | Minimum Essential Media (MEM) - No Phenol Red | thermofisher | Cat. No. 51200-038 |  |
| Chemical compound, drug | Characterized Fetal Bovine Serum | hyclone | Cat. No. SH30071.03 |  |
| Chemical compound, drug | Penicillin- Streptomycin (5,000 U/mL) | thermofisher | Cat. No. 15070063 |  |
| Chemical compound, drug | N-2 Supplement (100X) | thermofisher | Cat. No. 17502048 |  |
| Chemical compound, drug | Donor Equine Serum | hyclone | Cat. No. SH30074.03 |  |
| Chemical compound, drug | Sodium Pyruvate (100 mM) | thermofisher | Cat. No. 11360–070 |  |
| Chemical compound, drug | Neurobasal- A Medium | thermofisher | Cat. No. 10888–022 |  |
| Chemical compound, drug | B-27 Supplement (50x), serum free | thermofisher | Cat. No. 17504–044 |  |
| Chemical compound, drug | GlutaMAX supplement | thermofisher | Cat. No. 35050061 |  |
| Chemical compound, drug | Earle's Balanced Salts | sigmaaldrich | Cat. No. E3024 |  |
| Chemical compound, drug | Corning Collagen I, Rat | FISHER SCIENTIFIC | Cat. No. 354236 |  |
| Chemical compound, drug | Cover Glass | FISHER SCIENTIFIC | Cat. No. 12-545-80 |  |
| Chemical compound, drug | Papain | Worthington Biochemical | Cat. No. LS003126 |  |
| Chemical compound, drug | PDL (poly-D-lysine) | BD Biosciences | Cat. No. 40210 |  |
| Chemical compound, drug | Trypsin-EDTA (0.05%), phenol red | thermofisher | Cat. No. 25300–054 |  |
| Biological sample<br>(Rattus norvegicus, Female) | Sprague-Dawley<br>Timed-Pregnant rat,<br>E15 pups | Charles River<br>River |  | Pups of both genders |
| Software, algorithm | MATLAB | MathWorks | RRID:SCR_001622 |  |
| Software, algorithm | u-track2.0 | Jaqaman, K., et al. (2008), Nat.Meth. 5, 695-702 | Gaudenz Danuser Lab |  |
| Software, algorithm | ImageJ | https://imagej.nih.gov/ij/ | RRID:SCR_003070 |  |
| Software, algorithm | Fiji | http://fiji.sc | RRID:SCR_002285 |  |
| Software, algorithm | inkscape |  | RRID:SCR_014479 |  |
| Other | VGluT1-pHluorin | Drs. Robert<br>Edwards and<br>Susan Voglmaier<br>(UCSF) |  | Genetically<br>encoded optical<br>indicator of vesicle<br>release and recycling |

